# Evaluation of Deep Learning for predicting rice traits using structural and single-nucleotide genomic variants

**DOI:** 10.1101/2024.01.18.576088

**Authors:** Ioanna-Theoni Vourlaki, Sebastián E. Ramos-Onsins, Miguel Pérez-Enciso, Raúl Castanera

**Affiliations:** Centre for Research in Agricultural Genomics CSIC-IRTA-UAB-UB, Campus UAB, Edifici CRAG, Bellaterra, Barcelona 08193, Spain; Catalan Institute for Research and Advanced Studies (ICREA), Barcelona, Spain; Universitat Autónoma de Barcelona, Barcelona, 08193, Spain

## Abstract

Structural variants (SVs) such as deletions, inversions, duplications, and Transposable Element (TE) Insertion Polymorphisms (TIPs) are prevalent in plant genomes and have played an important role in evolution and domestication, as they constitute a significant source of genomic and phenotypic variability. Nevertheless, most methods in quantitative genetics focusing on crop improvement, such as genomic prediction, consider Single Nucleotide Polymorphisms (SNPs) as the only type of genetic marker. Here, we used rice to investigate whether combining the structural and nucleotide genome-wide variation can improve prediction ability of traits when compared to using only SNPs. Moreover, we also examine the potential advantage of Deep Learning (DL) networks over Bayesian Linear models, which have been widely applied in genomic prediction. Specifically, the performance of BayesC and a Bayesian Reproducible Kernel Hilbert space regressions were compared to two different DL architectures, the Multilayer Perceptron, and the Convolution Neural Network. We further explore their prediction ability by using various marker input strategies and found that exploiting structural and nucleotide variation improves prediction ability on complex traits in rice. Also, DL models outperformed Bayesian models in 75% of the studied cases. Finally, DL systematically improved prediction ability of binary traits against the Bayesian models.

## Introduction

Rice (*Oryza sativa*) constitutes a fundamental staple crop, essential to humans for its nutritional and caloric value. World rice production has reached a plateau <u>(FAO, 2023)</u>, and following the conventional breeding techniques, rice yield will soon not meet the high demand caused by the increasing world population. Therefore, we need to explore new approaches to secure global nutritional requirements by increasing at the same time the quality and quantity of rice yield. Genomic Prediction (GP) can help to achieve these goals, accelerating the breeding progress (<u>Meuwissen et al. 2001</u>). Various studies in plants have shown the effectiveness of GP in increasing breeding speed (<u>Jighly et al. 2019</u>, <u>Tessema et al. 2020</u>, <u>Xu et al. 2020</u>, <u>Krishnappa et al. 2021</u>). GP framework has been widely used in rice studies for predicting various quantitative traits, reporting moderate to high predictive performance (<u>Xu et al. 2021</u>). Complex traits are controlled by numerous loci that are difficult to detect with genetic mapping. GP assumes that quantitative trait loci (QTL) will be in linkage disequilibrium (LD) with molecular markers. Thus, instead of detecting all the QTL associated with a trait, an indirect association between marker and trait is exploited.

Conceptually, since the number of genotyped individuals *n,* is typically smaller than the number of molecular markers *p,* GP faces statistical challenges such as large sampling variance and increased mean-square error. To overcome this limitation, variables must be selected or restrictions on the solutions must be applied or sometimes both. The main classes of GP methods are the genomic relationship-based method such as Genomic Best Linear Unbiased Prediction (GBLUP, <u>VanRaden, 2008</u>) and the SNP effect-based methods such as the Bayesian family (<u>Meuwissen et al. 2001</u>, <u>Habier et al. 2011</u>, <u>Pérez and De Los Campos, 2014</u>) and LASSO <u>(Tibshirani, 2011)</u>. Some Bayesian models do not necessarily assume homogeneity across marker effects. They perform variable selection and shrinkage on the effects simultaneously using priors other than Gaussian. BayesC is an example of this category assuming as a prior a normal distribution with constant variance while a fraction of marker has no effect (<u>Habier et al. 2011</u>). On the other hand, methods such as GBLUP involve restriction on the square of solutions (L2 norm), with the effect of the markers assuming to be normally distributed with equal variance.

Deep Learning (DL) networks are a collection of machine learning algorithms that have exhibited excellent performance in some prediction tasks (<u>Min et al. 2017</u>; <u>Pattanayak 2017</u>). The DL models are trained in such a way to find complex relationships between data. DL networks consist of multiple layers and interconnected nodes. Each layer uses as input the output of the previous layer to optimize the prediction or classification. Numerous DL architectures have been proposed, such as Multi-Layer Perceptron (MLP), Recurrent Neural Network (RNN) and Convolutional Neural Network (CNN, <u>LeCun et al. 2015</u>). DL has been around for decades but only recently started to be widely implemented because of the easy implementation framework provided by various online libraries (e.g https://keras.io/, https://pytorch.org/). The performance of the DL networks depends on the accuracy of hyperparameter choice, which is not an easy task and requires abundant computation resources (<u>Young et al. 2015</u>, <u>Chan et al. 2018</u>). For a review of DL tools applied to genomic prediction, see <u>(Pérez-Enciso and Zingaretti, 2019).</u>

Despite their features, various works have shown a performance of DL in genomic prediction comparable to linear models (<u>González-Recio et al. 2014</u>; <u>Ma et al. 2017</u>; <u>Bellot et al. 2018</u>; <u>Montesinos-López et al. 2018</u>). <u>Zingaretti et al. (2020)</u> did not find a considerable advantage of DL over linear models, except when epistasis component was important. <u>Ehret et al. (2015)</u> found non-relevant differences between a GBLUP and a MLP model. In a wheat study <u>(Ma et al. 2017)</u>, DL performed better than GBLUP when used to predict phenotypes from genotypes. Similarly, <u>Gianola et al. (2011)</u> found that MLP performed better than a Bayesian linear model in wheat. In another study in wheat, <u>Pérez-Rodríguez et al. (2012)</u> extensively compared the prediction performance of Radial Basis Function Neural Networks and Bayesian Regularizes Neural Networks against several linear models and semiparametric models such as Reproducible Kernel Hilbert Space (RKHS). The authors concluded that the non-linear models, such as DL, demonstrated a higher prediction ability than the linear models. Evaluating the potential of DL algorithms, <u>Sandhu et al. (2021)</u> showed higher prediction ability against the linear models in spring wheat. For an extensive review in GP using DL models see <u>Montesinos-López et al. (2021)</u>.

While genomic prediction of quantitative traits has been broadly studied, binary and ordinal traits are of great importance in plant breeding as well. Particularly, in rice, culm morphology and stay green related traits are important targets for genetic improvement (<u>Kamal et al. 2019</u>, <u>Chigira et al. 2023</u>, <u>Lee and Masclaux-Daubresse 2021</u>). However, the prediction ability of different types of traits depends on the statistical models used. Thus, many studies in Genomic Selection (GS) focus on improving existing models or develop new ones as it was pointed out by <u>Montesinos-López et al. (2019)</u>.

Most of the studies in GP rely only on SNPs, disregarding other sources of genomic variation such as structural variants (SVs). Among the different types of SVs, TIPs account for an important fraction and play a key role in plant evolution, from domestication to adaptation and breeding <u>(Dubin et al. 2018)</u>. Studies in tomato and in rice found that the use of TIPs can lead to the identification of novel associations not detected by SNPs in genome-wide association studies (<u>Akakpo et al. 2020</u>; <u>Carpentier et al. 2019</u>; <u>Domínguez et al. 2020</u>; <u>Castanera et al. 2021</u>). In a recent study, we showed that TIPs explain a sizable fraction of the genetic variance in several rice agronomic traits and significantly improve genomic prediction (<u>Vourlaki et al. (2022)</u>. However, TIPs are not the only type of structural variation in the genome. Other SVs such as deletions, inversions, and duplications account for an important fraction of genetic variation and have been key in the domestication and diversification of plant crops <u>(Lye and Purugganan, 2019)</u>. SVs have a high potential to generate large effect mutations when they affect genes or gene regulatory regions. Over the last few years, several studies have demonstrated the importance of presence-absence variation and structural variation as a source of gene expression and phenotypic variability in plant crops including rice, tomato or soybean (<u>Qin et al. 2021, Alonge et al. 2020, Liu et al. 2020</u>).

Here we investigate whether merging all the structural and nucleotide genome-wide variation can improve phenotypic prediction compared to using only SNPs in rice. Finally, we further explore the performance of DL in GP by (i) using multiple marker input strategies, (ii) proposing several approaches to accommodate large scale marker information, and (iii) optimizing network architectures. We also provide documented python code based in tensorflow 2 <u>(Abadi et al. 2016) in the following link</u> https://github.com/ivourlaki/Deep-Learning-in-rice-for-prediction.git.

## Materials and Methods

### Rice accessions and traits

In this study we used a subset of 738 accessions from the 3,000-rice genome project <u>(Li et al. 2014)</u>. Accessions were chosen based on the availability of at least 15x of sequencing depth. The 738 accessions are representative of the main rice population groups: Aus/Boro (AUS, N=75), Indica (IND, N=451), Japonica (Jap,N=166), Aromatic (ARO, N=17). An additional group of admixed varieties (ADM, N=29) consisting of accessions that cannot be assigned to a specific rice group was also used. SNP-based group assignment from <u>Alexandrov et al. (2015)</u> and <u>Sun et al. (2017)</u> was used to identify the different subsets of this study. Studied traits are publicly available at IRRI SNP-Seek database (https://snp-seek.irri.org/). Among continuous traits, grain weight and time to flowering were used, whereas culm diameter (1^st^ internode) and leaf senescence were selected among the binary traits available. Some of the ordinal traits were binned to balance the number of observations per class. Particularly, in leaf senescence values equal to 1 were assigned to class 1 while the rest ones range from 2-9 were recorded as 2 and were assigned to class 2. Finally, time to flowering was log-transformed.

### Markers

We used the filtered SNP dataset in <u>Vourlaki et al. (2022)</u>. Specifically, a binary ped file format with the 3K RG CoreSNPs dataset for all chromosomes was downloaded from the SNP-Seek database. The original dataset consisted of 404,399 bi-allelic SNPs from 3,034 rice accessions, including the 738 accessions selected. We filtered out markers with minor allele frequency ≤ 0.01 and missing rate > 1% using plink2 (<u>Purcell et al. 2007</u>; <u>Chang et al. 2015</u>). Also, missing genotypes were imputed using Beagle 5.2 with default parameters (<u>Browning et al. 2018</u>). After filtering the final dataset consisted of 228,871 SNPs.

The TIP dataset described in <u>Castanera et al. (2021)</u> (containing two categories: MITE-DTX and RLX-RIX) was complemented with non-TE deletions (DEL), duplications (DUP) and inversions (INV) downloaded from SNP-seek database (3K RG Large Structural Variants release 1.0). We filtered out SVs events containing multiple overlapping deletions as these complex variants are difficult to genotype with short reads and are thus less reliable. SVs genotypes were recorded as 0/1 (absence/presence). All the markers with minor allele frequency ≤ 0.01 were filtered out. Finally, the dataset used in our analysis consists of 52,120 MITE-DTX, 21,517 RLX-RIX, 74,136 DEL, 25,670 DUP and 7,527 INV.

### Genetic variance inference

To estimate the genetic variance components explained by each marker set, we fitted the following linear model using RKHS <u>(Gianola et al. 2006)</u>:

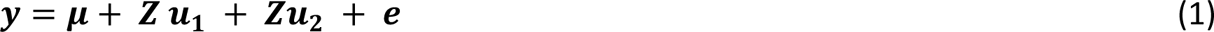

where *μ* is the general mean, *y* is the phenotype vector of size n (the number of accessions), *Z* is an identity incidence matrix, *u*_1_and *u*_2_ are random effects of each of the marker groups and e is the residual. Random effects are assumed to be normally distributed 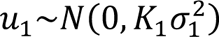, 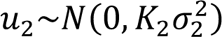, with constant variance 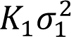 and 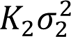. Where *K*_1_, *K*_2_ are genomic relationship matrices (GRM) obtained from the markers used in the corresponding model. We fitted the model five times using as *K*_1_ the GRM form SNPs while as *K*_2_, GRM was obtained from MITE-DTX, RLX-RIX, DEL, DUP, INV, separately. The GRM were calculated using AGHMatrix <u>(Amadeu et al. 2016)</u>. Model was implemented in BGLR package <u>(Pérez and de Los Campos 2014)</u> using default priors to estimate 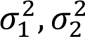.

### Genomic Prediction Models

#### Bayesian Regression Models

Two Bayesian methods were employed in this study: Bayesian RKHS and BayesC. RKHS is a kernel-based method that uses a ridge regression L2 regularization technique like GBLUP. BayesC is a variable selection method that estimates the effect of the markers. Both methods were applied to each trait separately. Particularly, for each method, various models were designed and applied comparing the predictive performance of using all the markers together versus using only SNPs. For RKHS, the models are described as follows:

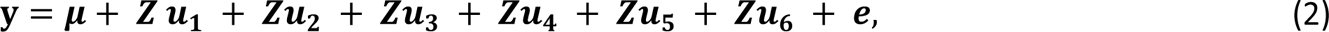

Where *u*_1_, *u*_2_, *u*_3_, *u*_4_, *u*_5_, *u*_6_ are random effects *u*∼*N*(0, *Kσ*^2^) with *K* corresponds to a GRM computed using SNPs, MITE-DTX, RLX-RIX, DEL, DUP, INV, respectively. Apart from model (2) where the different marker types are applied at the same time, other models were studied as well. Particularly, we were interested in investigating whether one matrix represented by different type of markers could result in a higher prediction in comparison to model (2) and to use only SNPs. As a result, the following models were studied:

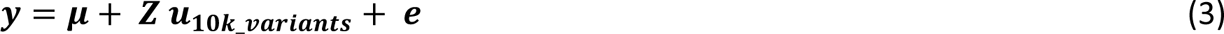

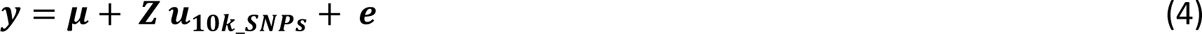

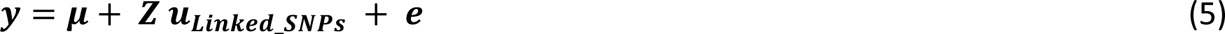

Where *u*_10*k*_*variants*_, *u*_10*k*_*SNPs*_, *u_Linked_*__*SNPs*_, are the effects of the 10,000 most associated markers among the six sets, the 10,000 most associated SNP effects and by using the SNPs linked to structural variation. LD between SNPs and SVs was calculated using ngsLD software <u>(Fox et al. 2019)</u>, and we considered that a SNP was linked to an SV when r^2^ >= 0.8.

For BayesC, the complete models were:

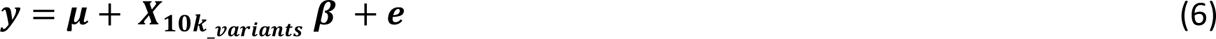

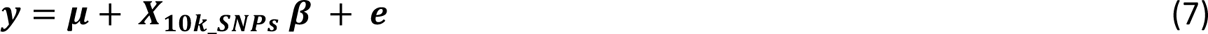

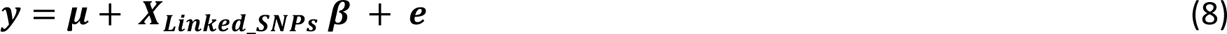

Where *X*_10*k*_*variants*_, *X*_10*k*_*SNPs*_, *X_Linked_*__*SNPs*_ are the standardized genotypes for the 10,000 most associated markers among all marker sets, the 10,000 most associated SNPs and the linked SNPs respectively. The 10,000 most associated were selected based on a linear regression-based genome-wide association analysis (GWAS) analysis. A detailed description is given in “Input Strategies” section. Finally, where *β*, is the vector of effects for the corresponding matrix.

Using either RKHS or BayesC, phenotypes to be predicted were removed from the dataset and the model fitted using the remaining phenotypes. Prediction ability was assessed by computing two different metrics related to the type of trait. We computed the mean squared error (MSE) between predicted and observed phenotypes for the quantitative traits, whereas binary cross-entropy was employed for the binary traits. Both models were implemented using BGLR package. BayesC assumes that a proportion of markers will have zero effect with probability sampled from a beta distribution, *π*∼*Beta*(*p*_0_, *π*_0_). Here we chose *p*_0_= 5 and π_0_ = 0.01. For the case of binary traits option “response_type=ordinal” was applied in both methods (RKHS, BayesC). Finally, BGLR was run for 100,000 iterations using default priors for RKHS.

#### Multilayer Perceptron

One of the most popular DL architectures is the Multilayer Perceptron (MLP). MLP is a fully connected feedforward artificial neural network which transforms any input dimension to the desired dimension. All the neurons are connected to every neuron in the previous layer and then connected to every neuron in the next layer. Each neuron receives the initial inputs multiplied by a corresponding weight coefficient. Then the sum of all inputs multiplied by weight plus a bias, is passed to an activation function which introduces the non-linearity to the network transforming the inputs accordingly. We can represent the output of each hidden layer as (note the transposes):

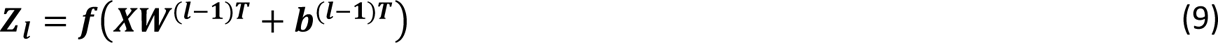

Where *Z_l_* is the output of the l layer, *b*^(*l*-1)*T*^ is the bias vector of the first layer, *X* is a single matrix of all training examples so that we could compute all the prediction using a single matrix multiplication, *W*^(*l*-1)*T*^is the weight matrix and *f* is a nonlinear activation function. The model is trained successively, that is, the output of neurons from the previous layer will be the input for the next layer. Figure 1 shows the basic workflow of MLP network.

**Figure 1:**
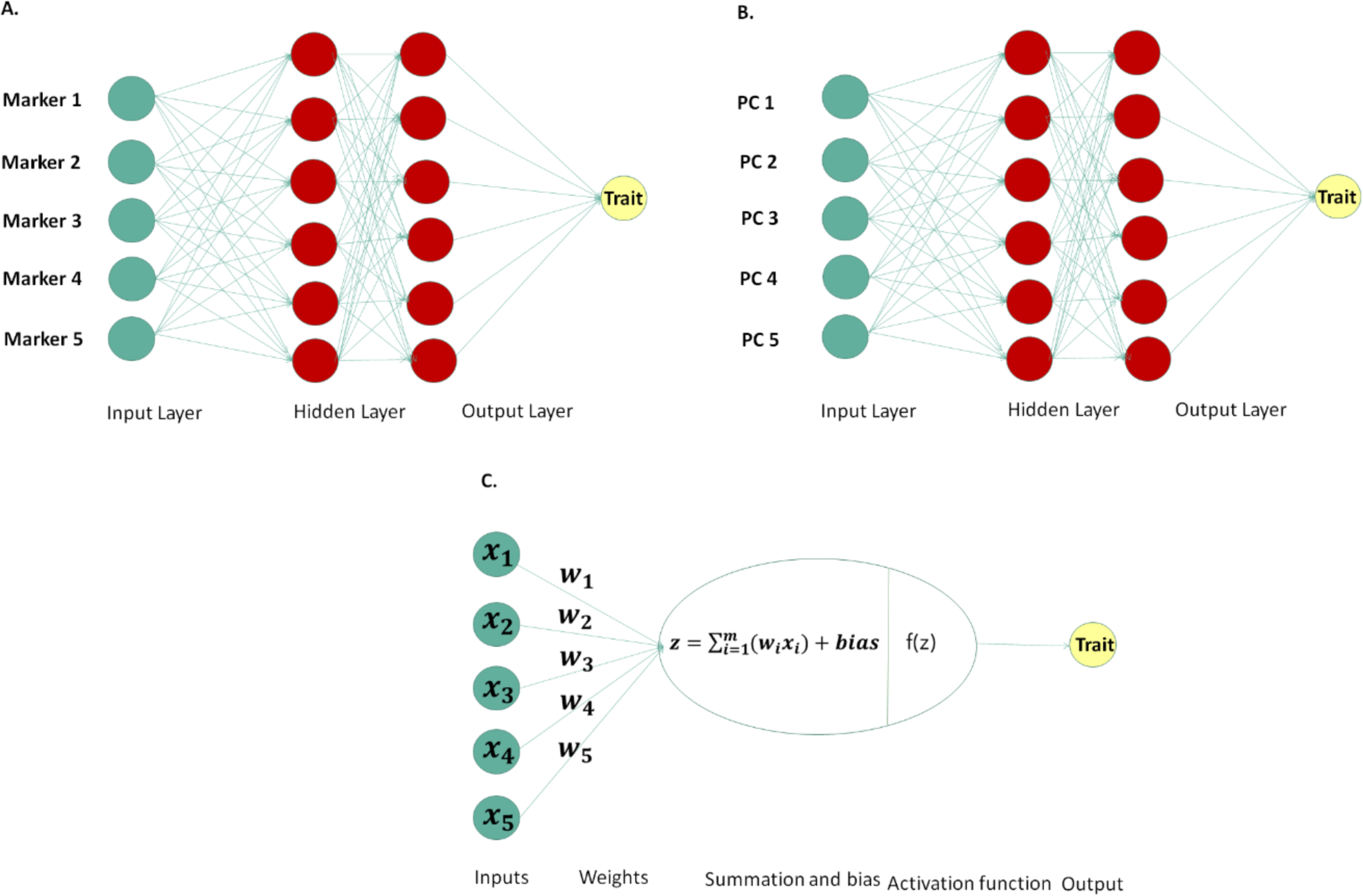
Multilayer perceptron (MLP) representation with markers (A) and PCs (B) as input layers; bottom center shows the basic workflow of a perceptron or else neuron (C).

#### Convolutional Neural Networks

Convolutional neural networks (CNNs) can utilize spatial relationships between nearby variables (e.g., pixels) of the input matrix. This architecture can accommodate situation where input variables are distributed along a space and are associated with each other such as linkage disequilibrium between nearby markers <u>(Pérez-Enciso and Zingaretti, 2019)</u>. A CNN has hidden layers which typically consist of convolutional layers, pooling layers, flatten layers and fully connected dense layers. In each convolutional layer, CNN automatically performs a convolution that is a linear operation performed along the input of predefined width and strides by applying kernels or filters. The weights used are the same for all marker windows. The filter moves along windows of same sizes consist of markers performing a multiplication operation (dot product) until the entire matrix is traversed. The output of the convolutional function can be described as an integral transformation <u>(Widder, 1954)</u>, as follows:

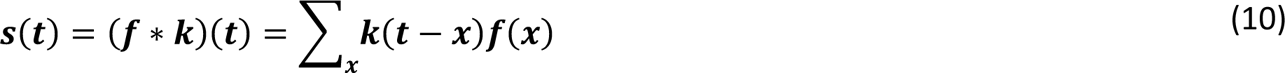

where *k* represents the kernel, convolution is the transformation of *f* into *s*(*t*). The operation is performed over an infinite number of copies *f* resulting in the weighted sum shifting over the kernel. An activation function is applied after each convolution to produce the output layer. After nonlinearity has been applied to the feature map produced by the first layer, a pooling layer usually follows, aiming to reduce the dimensionality and smoothen the representation (Figure 2). The benefit of using CNN is their ability to develop an internal representation of a two-dimensional matrix extracting the most important features. CNN leverages the fact that nearby input variables are more strongly related than the distant ones.

**Figure 2:**
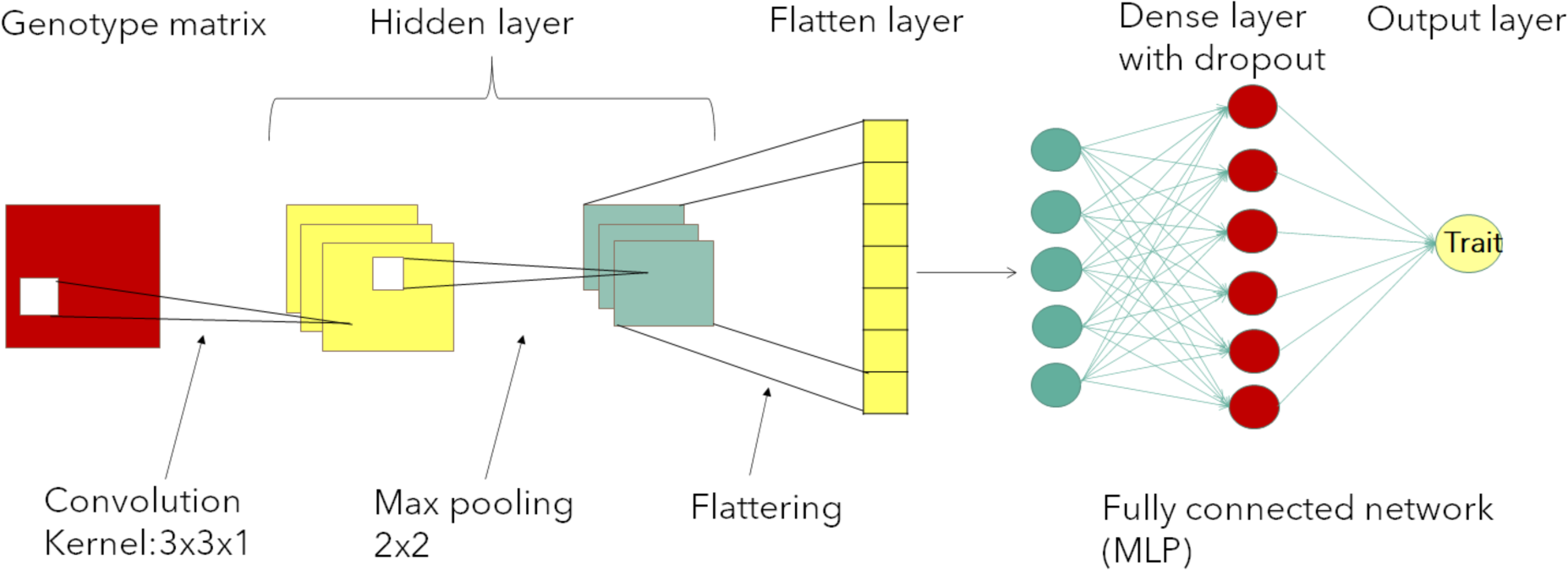
Convolutional Neural Network (CNN) representation used in study.

### Cross-validation and Independent Prediction

Here we evaluate the prediction accuracy by following two challenging validation scenarios relevant for breeding programs: prediction of individuals from two different population groups and prediction of randomly selected individuals from the rest ones. For the first strategy, we predicted performance of the admixed (ADM, N=29) and aromatic (ARO, N=17) groups using the rest accessions (IND, JAP and AUS/Boro). Since accessions to be predicted are not phylogenetically close to the accessions in the training set, it would be expected a low prediction ability from the models for this scenario.

In the second strategy, prediction accuracy was evaluated by implementing a 10-fold cross-validation (CV) where training population consisted of 90% of the data and testing set included 10% of the remaining data. Analysis was performed in each of the ten training sets separately assuming ten different validation scenarios. Since accessions are randomly selected and not based on their population group, samples in the training set might be related to the predicted ones. Note that, in the case of DL application, training population was further split in a validation dataset which included 20% of the training data set (Figure 3).

**Figure 3:**
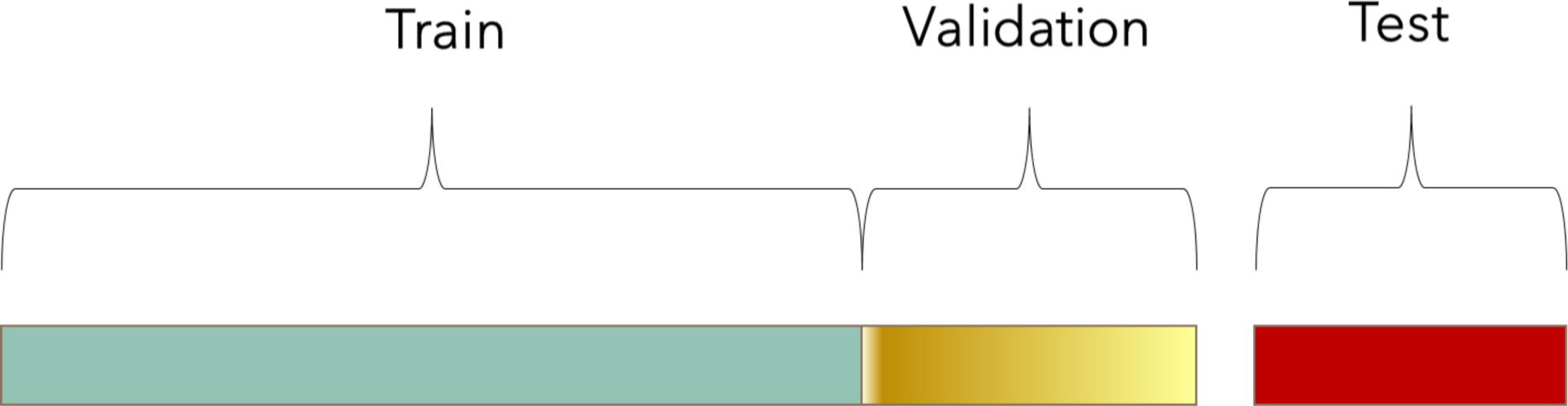
A visualization of how the three data sets, training, validation, and test are divided.

Validation data set is used during the training process of our network to provide an unbiased evaluation of a model fit on the training dataset while tuning model hyperparameters. It is important to mention that the model “sees” the data and use it for an evaluation of the process but never “learn from these”. After the model was trained, we retrieved the best hyperparameters and performed prediction using the test dataset. The test dataset provides the gold standard used to make an unbiased final evaluation of the model. It is used only once a model is completely trained using the train and validation sets.

### Marker Input strategies

We used different marker input strategies aiming to enhance network flexibility and thus improve prediction ability. Four strategies were designed as follows:

1. Most associated markers: In the first input strategy, we merged the structural and nucleotide genomic-wide variation to test whether prediction accuracy can be improved. However, using the whole six genotype matrices (SNPs, MITE-DTX, RLX-RIX, DEL, DUP, INV) would add a high complexity in our network that might cause an overfitting. Studies have shown that using a subset of markers can result in equivalent prediction ability to using all the data sets (<u>Müller et al. 2017, Bellot et al. 2018</u>, <u>Vourlaki et al. 2022</u>). Thus, from the total of 409,892 molecular markers we selected the 10,000 most associated to the traits of interest. Specifically, we performed a genome wide association study (GWAS) fitting a linear model (single-marker regression analyses) to find associations between each of the six-marker set and each of the four traits (4×6). For each fitted model, a p-value corresponding to each marker was collected. From the collection of the p-values the 10,000 most associated was selected. GWAS was performed only to the training sets. Since we followed two different cross-validation strategies the process was repeated for each of those, that is for the across population training set, for the ten partitions training sets and for each trait. This strategy (hereafter referred to as “COMBINED_VARIANTS”) was applied to DL and Bayesian linear models. Additionally, we selected the 10,000 most associated SNPs (hereafter referred as “SNPs”) to perform the same analysis and compare directly to the COMBINED_VARIANTS strategy.
2. Linked SNPs: Using causal variants associated to the specific traits can result to prediction accuracy almost 1 <u>(Pérez-Enciso et al. 2015)</u>, and structural variants are often causative of trait variability. Nevertheless, SNPs are easier to genotype in populations than SVs (ie, by genotyping chips). We reasoned that SNPs in LD with SVs could be used as SV replacement and would be easier to use in further experimental or breeding programs. Hence, we used ngdLD to detect SNPs in LD with SVs (r2 >= 0.8) and used them for prediction. The software was applied to each trait and cross-validation scenario using only the training set. Then, for each case the unique pairs of SNPs-SVs with LD >= 0.8 were selected. The type, position and chromosome of the variants meeting the criteria were collected in a list and is provided in the github repository. The linked SNPs sets were used as marker sets across all the analysis. The selected abbreviation for this input strategy was “LINKED_SNPS”. The average number of linked SNPs across the analyses was ∼ 12,000.
3. PCs single matrix: In the third strategy, we exploited the advantages of principal components analysis (PCA) by incorporating it to neural networks. Studies have shown that using principal components in DL framework can be particularly advantageous <u>(Seuret et al. 2017)</u>. In our study, PCs were computed based on eigenvectors for each of the obtained GRM. We run the analysis by feeding the network with a single matrix merging the PCs computed by the six GRMs introducing them as a single layer (Figure 1 (B)). Also, a separate analysis was performed for SNPs and linked SNPs testing whether this strategy will enhance the performance of the model. The selected abbreviation for this input strategy was “COMBINED_VARIANTS” with method applied MLP_PCs.
4. Multiple Inputs: Here, we tested whether multiple input strategy could improve the prediction of traits. Particularly, we used the six PCs sets (computed by the GRM of each marker set) as six inputs feeding to the network in different input layers simultaneously. Other works have shown that a multiple input strategy can reduce overfitting and computational cost while at the same time exploits mixed data improving prediction (<u>Livieris et al. 2020</u>, <u>Xiong et al. 2021</u>). Thus, the network accepted six different input layers which independently forwards in six different hidden dense layers. Next the six layers are merged by a concatenate layer (Figure 4). In order to compare directly this DL strategy against Bayesian Linear models, RKHS was performed with model eq. 2 using all the GRM as inputs at the same time. The selected abbreviation for this input strategy was “MULTIPLE INPUTS”.

**Figure 4:**
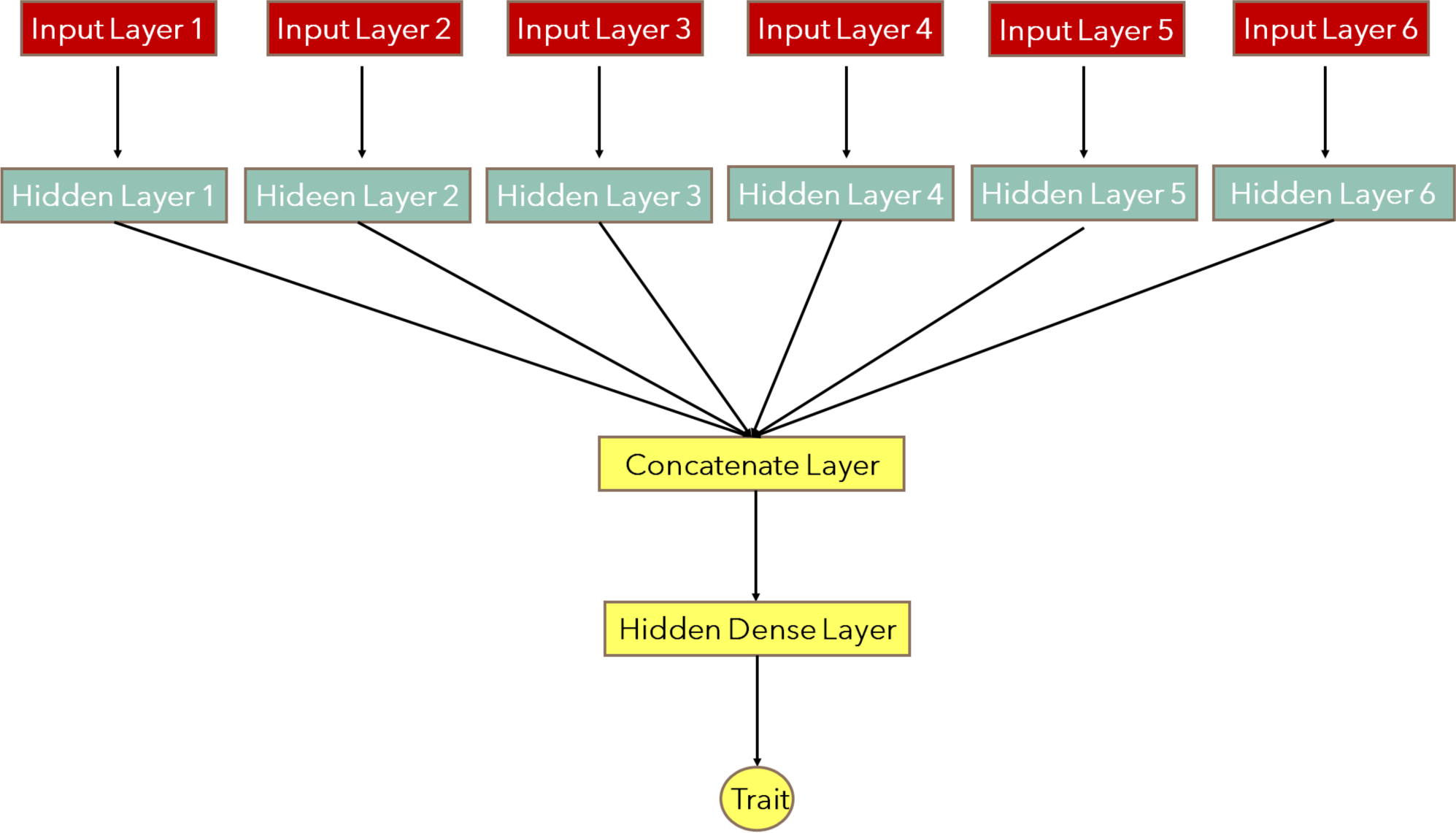
Representation of Multiple inputs strategy employed in the present study.

### Optimization of Hyperparameters

CNNs, MLP, BayesC and RKHS were implemented using the COMBINED_VARIANTS (SNPs+SVs), as well as SNPs and LINKED_SNPs as inputs separately. Additionally, MLP network was employed for using the six PCs as a single input matrix, as six different input layers and for PCs produced by GRM of SNPs. All the models were applied separately to each trait and to eleven cross-validation scenarios (10-fold and ARO/ADM across-population prediction). Table 1 shows the different models implemented in our analysis.

**Table 1:** Summary of the analyses performed.

| Trait | MLP, CNN, BayesC, RKHS, MLP PCs |  |  | RKHS, MLP PCs |  |
| --- | --- | --- | --- | --- | --- |
|  | Combined variants | SNPs | Linked SNPs | Multiple Inputs |  |
| Culm diameter | X | X | X | X | X |
| Leaf senescence | X | X | X | X | X |
| Grain weight | X | X | X | X | X |
| Time to flowering | X | X | X | X | X |

For each of the different runs, hyperparameter tuning was performed obtaining the best hyperparameters and then retrained the model with the hyperparameters obtained by the search. Here Keras Tuner <u>(O’Malley et al. 2019)</u> library was used to pick the optimal set. Hyperparameters are the variables that control the training process and the topology of our model. When the model is built for hyperparameter tuning, the search space is also defined in addition to the model architecture. Then a tuner must be selected to determine which hyperparameter combinations should be tested. In our analysis we used the Hyperband tuner. The Hyperband tuning algorithm uses adaptive resource allocation and early stopping to quickly converge on a high-performing model. The algorithm trains a large number of configurations for a few epochs and carries forward only the top-performing half of models to the next round <u>(Li et al. 2018)</u> evaluating the performance by computing the MSE (for quantitative traits) and binary cross-entropy (for binary traits) were computed on a held-out validation set. The best model is the one that minimizes errors. After the hyperparameter search was finished, we evaluated the model on the test data and performed prediction computing the pre-mentioned evaluation metrics of interest on the test data set. Figure 5 displays the suggested scheme.

**Figure 5:**
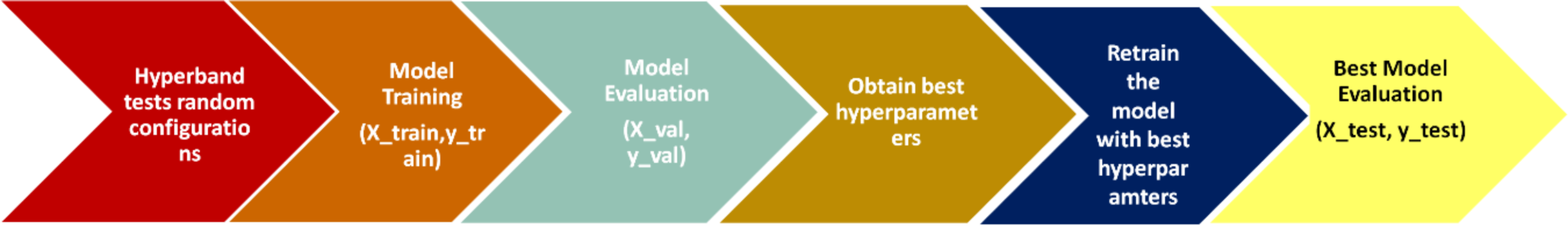
Figure depicts the basic scheme performing from hyperband tuner to determine the best configuration towards to the final evaluation of the model.

DL performance is controlled by various parameters and thus the optimization of the hyperparameters is not a trivial step. Here, we designed the tuner search space based on the available literature <u>(Sandhu et al. 2021</u>, <u>Zingaretti et al. 2020)</u>. The hyperparameters chosen to be optimized were: activation function (Relu, Tanh, Linear), number of neurons for the first layer in MLP and number of filters in CNN (16,38,64,128), number of hidden dense layers for MLP (0,1,2,3) while for CNN (1,2,3), number of neurons of hidden layers (2,4,8,16), numbers of optimizers (Adam, RMSprop, SGD), dropout rate (0,0.05,0.1,0.15,0.2,0.25,0.3), L1 and L2 regularizers with optimized weight decay parameter (0.001, 0.01, 0.1). For the output layer we used one unit with linear as activation function for the quantitative traits whereas sigmoid was used for the binary traits as it is suggested <u>(Montesions-López et al. 2022)</u> For the hyperparameter optimization, 80% of training set was used and the remaining 20% validation data set was applied for inner testing. Training a DL network that can generalize well new data set is a challenging issue. A model with too little capacity cannot learn from the data, a problem known as underfitting, whereas a model with a large capacity can learn and fit too well to the training dataset results in overfitting. For avoiding and reducing the effects of these two phenomena there are techniques that can be adjusted to a DL network.

Here we used two regularization techniques such as L1 and L2 with a weight decay parameter. These techniques penalize the weight values of the network making values tend to zero and negative equal to 0 and avoid a parsimonious model. L1 adds “squared absolute value of magnitude” of coefficient as penalty term to the loss function while L2 adds “squared magnitude” of coefficient as penalty term to the loss function. We added L1 and L2 regularizers in the hidden layers. Additional to the regularization, dropout and early stopping were applied to reduce the effect of overfitting and underfitting on our models. A dropout layer was applied before the output layer. Our analysis was implemented using Tensor Flow 2.8.0 library with Keras 2.8.0 interface and Keras Tuner 1.1.2.

## RESULTS

### Phenotypic Structure and Genetic Inference

We used PCA to determine the underlying structure of our data and the direction of the maximum variation when projected in a lower dimension space. Figure 6 shows the projections of variables of each trait onto the principal components. The length of the arrow is proportional to trait contribution, whereas the angle between arrows shows whether traits are correlated (pointed out in the same direction) or not. An analysis in two principal components showed that the first component depends on grain weight, which contributes the most to the total phenotypic variation. The main contributors to the second component in descending sequence are Time to flowering, Culm Diameter and Leaf Senescence.

**Figure 6:**
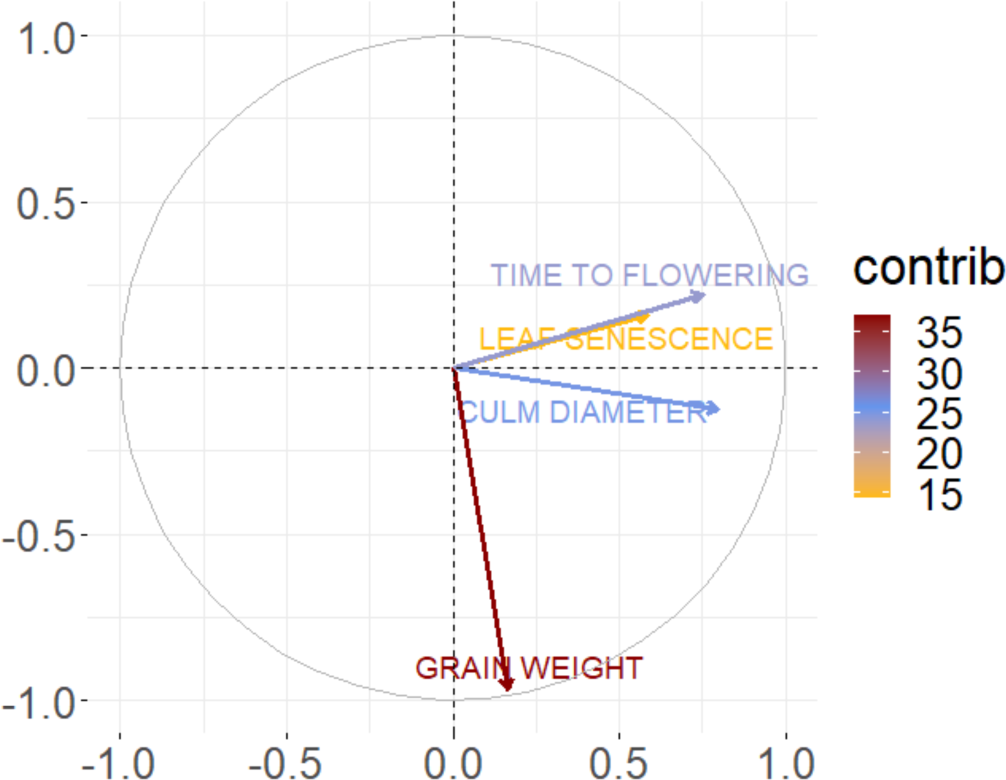
PCA loadings of each trait for the two first standardized principal components. The different colors indicate the percentage of contribution to the PCAs displayed as “contrib”.

Genetic variance estimates were obtained for each trait (Figure 7). Particularly, we estimated the genetic variance explained by each SV marker set in comparison to SNPs in order to understand the relative importance of each set to determine the observed phenotype. Figure 7 shows that structural variants explain a significant fraction of genetic variance, larger than that explained by SNPs in the two binary traits, Culm diameter and Leaf Senescence. Among the different types of SVs, TE-related variants (MITE-DTX and RLX-RIX) explained more genetic variance than non-TE variants. Among the latter, deletions explained more variance than duplications and inversions.

**Figure 7:**
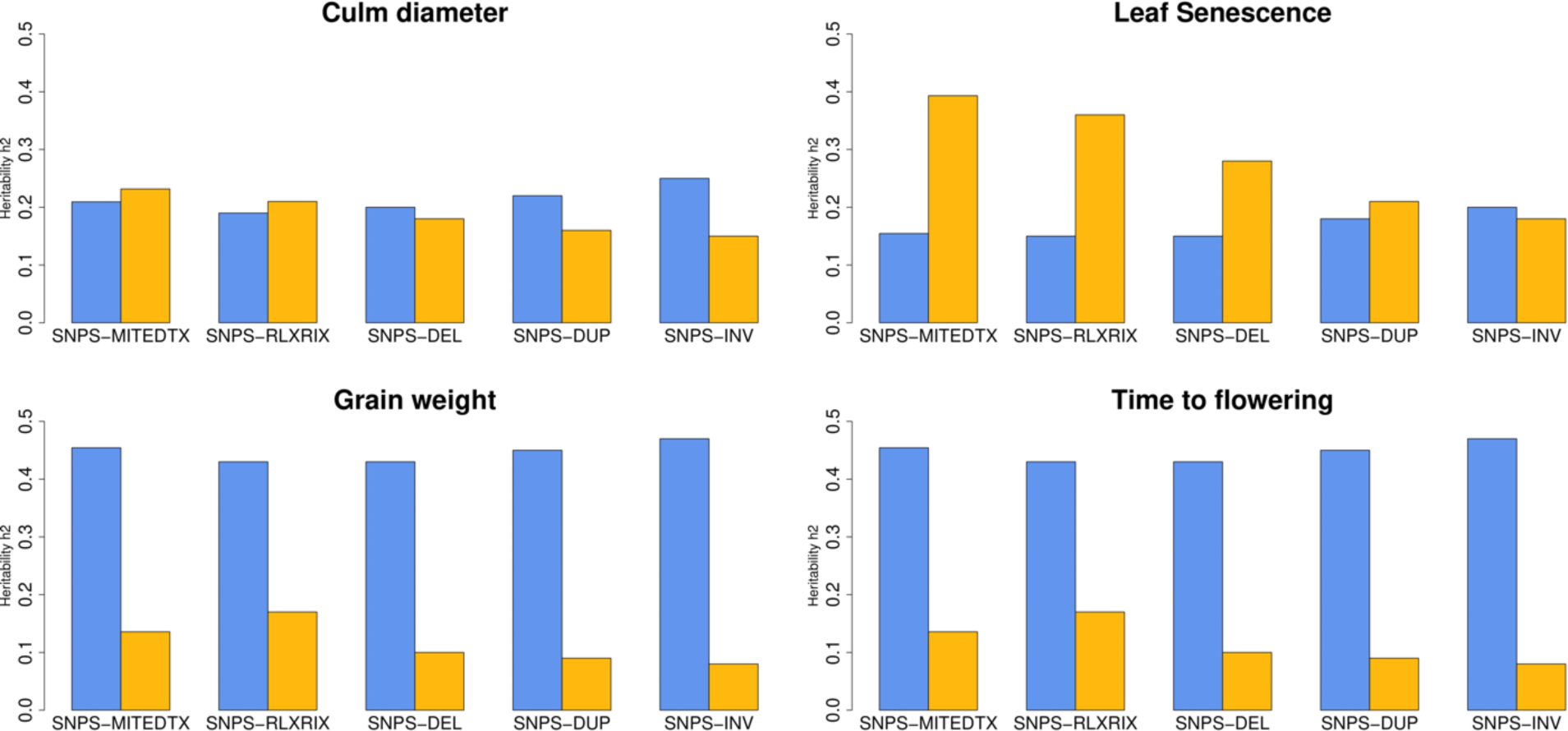
Means of posterior distributions of genetic variances explained by each marker set. Blue bars represent SNPs and yellow bars are the different kinds of SVs.

### Comparison of Model performances

The prediction ability of DL implementations was compared to those of Bayesian regression using RKHS and BayesC for each trait and under eleven cross validation sets. Particularly, we assessed prediction by following two different validation strategies, prediction using ten randomly selected training sets produced by a 10-fold cross validation strategy and prediction across populations. All the models were applied separately to each of the eleven validation scenarios (see Materials and Methods). Figure 8 shows the performance of each of the models under the 10-fold cross validation. We used binary-cross entropy as evaluation metric for binary traits and MSE for quantitative traits. Other metrics can also be applied (e.g Pearson’s correlation for quantitative traits, accuracy for binary traits) however here we measure the prediction ability in terms of loss of the model as previously reported <u>(Montesinos-López et al. 2023)</u>. The highest prediction ability (minimum loss) for culm diameter was obtained using CNN and linked SNPs (loss = 0.58). For leaf senescence the best prediction value was reported using MLP with PCs computed by the combined variants (Loss = 0.576). For the case of quantitative traits, Bayesian Regression models showed higher prediction accuracy values than those of DL models. Particularly, grain weight and time to flowering was better predicted under RKHS model using multiple inputs (Loss = 0.72 and 0.33, respectively).

**Figure 8:**
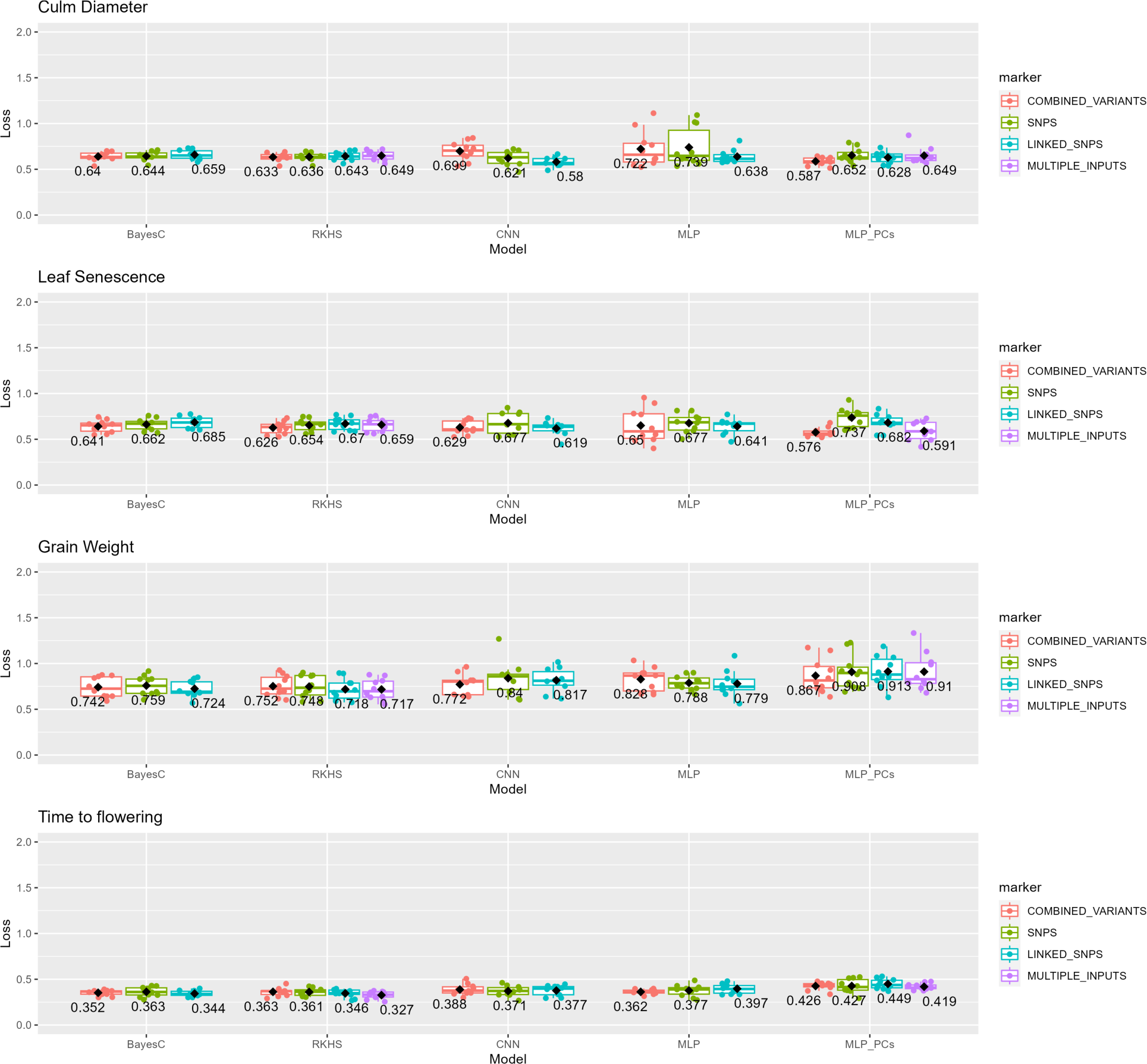
Performance of each of the model-input combinations under the 10-fold cross validation strategy and four predicted phenotypes. Each model was applied separately to each of the ten partitions. Points represent the evaluation metric for each partition whereas the boxplot shows the distribution of the numerical values displaying the data quartiles. The value that appears in bold is the average value of each model. The y-axis shows the loss metric values which for binary traits is the binary-cross entropy and for quantitative traits the MSE.

In the independent prediction, phenotypes of all ADM and ARO accessions were predicted given the rest of the accessions. Figure 9 shows the prediction ability for across population strategy under eleven different models. Here, DL models outperformed Bayesian ones in all the traits. Particularly, culm diameter and leaf senescence were better predicted under the MLP_PCs model and “Combined_VARIANTS” marker strategy as in 10-partitions. However, grain weight and time to flowering showed the lowest loss values under CNN with SNPs (Loss = 0.96) and MLP with combined variants (Loss = 0.44) respectively. In general, time to flowering was better predicted compared to the rest traits in both cross-validation strategies. On average, prediction across populations was less accurate for the quantitative traits than in 10-fold scenarios as it was expected because of the more distantly related raining and test data sets.

**Figure 9:**
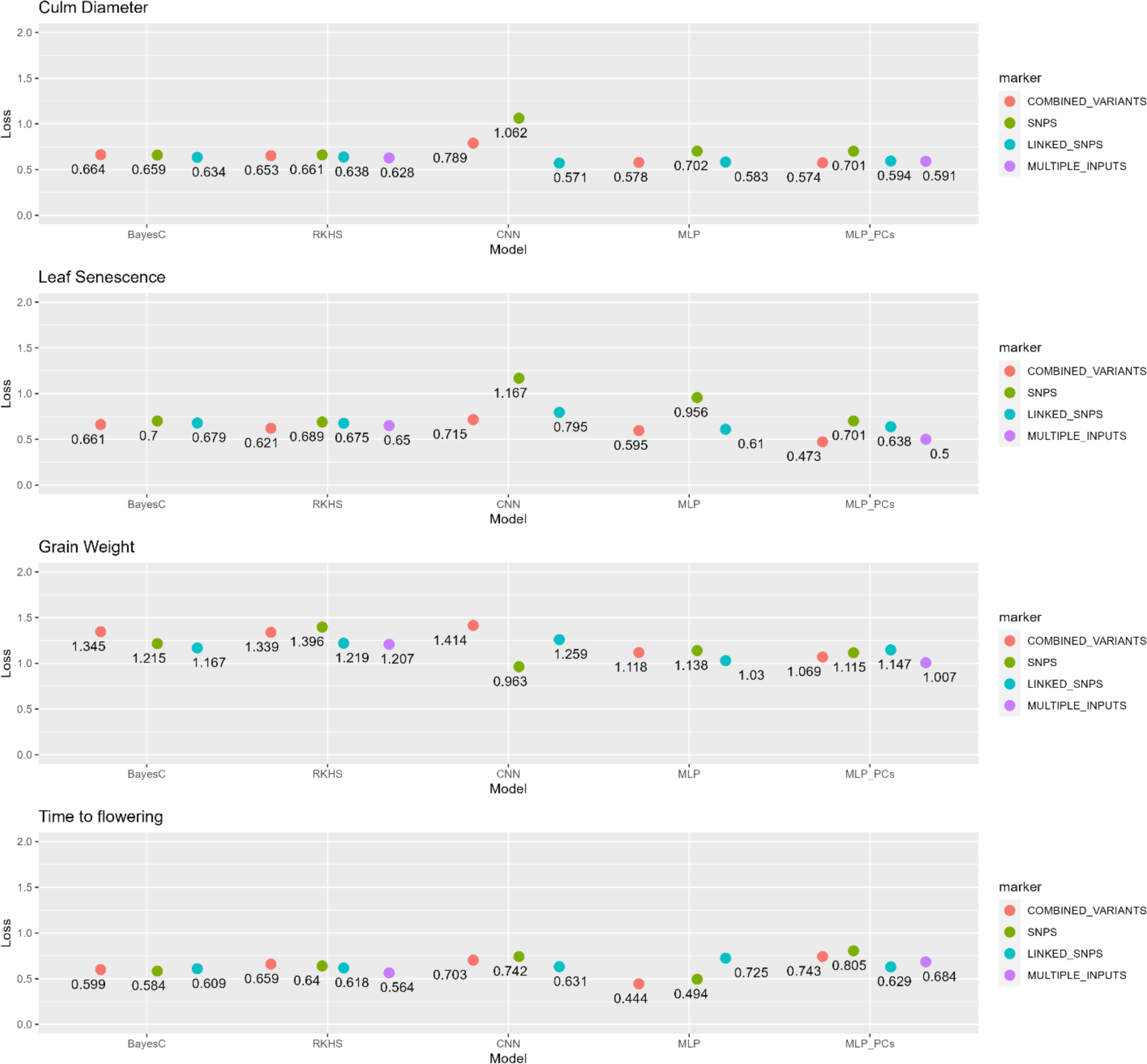
Performance of each of the model-input combinations under the across population strategy. Points represent the evaluation metric. The y-axis shows the loss metric values which for binary traits is the binary-cross entropy and for quantitative traits the MSE.

### Impact of marker selection on genomic prediction

Our results showed that phenotypic traits such as leaf senescence and time to flowering were better predicted using combined variants or multiple inputs. Also, using SNPs linked to SVs exhibited an efficient prediction ability especially for culm diameter under both validation strategies. We observed that incorporating structural variation in a genomic prediction framework either combining with SNPs or generating the linked SNPs to these variants resulted in an improved prediction performance in near 90% of the studied cases against using only SNPs (Table 6).

**Table 6:** Minimum prediction loss and corresponding model with input strategy.

| Traits | Validation Strategy |  |
| --- | --- | --- |
|  | 10-folds partitions | ARO/ADM accessions |
| **Culm diameter** | 0.580 (CNN, Linked SNPs) | 0.571 (CNN, Linked SNPs) |
| **Leaf senescence** | 0.576 (MLP_PCs, Combined variants) | 0.473 (MLP_PCs, Combined variants) |
| **Grain weight** | 0.717 (RKHS, Multiple inputs) | 0.963 (CNN SNPs) |
| **Time to flowering** | 0.327 (RKHS, Multiple Inputs) | 0.444 (MLP, Combined variants) |

## Discussion

In this study we investigated whether combining structural and nucleotide genome-wide variation for genomic prediction can improve prediction ability for important agronomic traits in rice. Previous studies on plants have shown the association between structural variants and phenotypic traits <u>(Żmieńko et al. 2014, Shang et al. 2022, Zhou et al. 2022)</u>, and some have been demonstrated to be the causal variants for a diversity of phenotypes across major traits in plants <u>(Sutton et al. 2007</u>, <u>Cook et al. 2012)</u>. In example, late or early flowering on wheat depends on the increased copy number of Vrn-A1 and Ppd-B1 genes respectively <u>(Würschum et al. 2015)</u>. In addition, plant height in wheat is associated with a specific tandem duplication <u>(Li et al. 2012)</u>. The strong regulatory potential of SVs could be an explanation for the high performance of SVs in the prediction of phenotypic traits. However, the incorporation of SVs in breeding programs demands their genotyping to be automatized. This can be a complex task, as SVs are highly diverse and commercial SV genotyping assays do not exist. Our results show that linked SNPs can be effectively used to indirectly incorporate SVs in genomic prediction. We propose that SNPs linked to know SV variants (ie, resulting from recent pangenome studies) constitute a promising marker resource to be used for future genomic prediction analyses.

A second objective addressed is the evaluation of the performance of DL and Bayesian methods to predict agronomically important traits in rice. Culm diameter, leaf senescence and time to flowering are correlated (Figure 6), whereas grain weight is uncorrelated to them. Traits such as time to flowering and grain weight are polygenic, controlled by many QTLs of large effects <u>(Begum et al. 2015</u>, <u>Xu et al. 2015</u>, <u>Chen et al. 2021)</u>. Studies in culm diameter have shown that it is controlled by at least twelve QTLs associated with lodging resistance in dry direct-seeded rice <u>(Yadav et al. 2017)</u>. In addition, delayed leaf senescence or stay-green is associated to forty-six QTLs that made up the genetic basis of this important trait in rice <u>(Jiang et al. 2004)</u>. Genomic prediction of traits such as time to flowering was quite accurate with the loss metric reported being the lowest values across all the study (average MSE value equal to 0.33, Table 6). For leaf senescence the GP ability was lower than that in time to flowering yet accurate. It is worth mentioning that, in the ARO/ADM validation strategy the prediction ability of leaf senescence and time to flowering was improved by DL against the best values of Bayesian models by 24% and 21% respectively (Figure 9). Since the genetic relatedness of the accessions used for training increases prediction accuracy, it is interesting that DL models outperform the Bayesian ones in both binary traits for genetically distant-lines. (Table 6).

Increasing the prediction accuracies of traits in rice breeding is challenging but at the same time of high importance, taking into consideration the increasing environmental constraints that limit world production. New methods attempt to improve prediction of agronomic traits promising lower computational cost and better results. DL is a state-of-the-art method applied in many different fields, and many recent studies have started to compare DL with standard linear models for genomic prediction <u>(González-Recio et al. 2014</u>; <u>Ma et al. 2017</u>; <u>Bellot et al. 2018</u>; <u>Montesinos-López et al. 2018</u>, <u>Zingaretti et al. 2020, Sandhu et al. 2021</u>, <u>Montesinos-López et al. 2019</u>, <u>Mon- tesinos-López et al. 2023</u>). Here, we studied the performance of DL models for predicting complex traits in rice comparing them to Bayesian regression methods under different input strategies and scenarios. Overall, our results showed that DL can increase prediction accuracy compared to Bayesian methods in 75% of the implementations. Across DL architectures, MLP and CNN were the optimal choices in the same number of cases depending on the trait and training population. This observation shows that there is not a clear winner, as evidenced by contrasting findings in the literature, where MLP outperforms CNN according to <u>(Sandhu et al. 2021)</u>, whereas (<u>Bellot et al. 2018</u>, <u>Zingaretti et al 2020</u>) report the opposite trend. For the case of Bayesian regression models, RKHS clearly outperformed BayesC.

Another critical and challenging issue in DL models is the optimization of hyperparameters, mainly due to the high computational cost. The tuning of the hyperparameters for each trait depends on the genetic basis and architecture of the trait. As we show in Supplementary Tables 1-4, different combinations of hyperparameters were selected for the various traits as the prediction ability is highly associated with the interaction of these factors <u>(Bellot et al. 2018</u>, <u>Montesinos-Lopez et al. 2018</u>). We observed that Tanh was the most useful activation function in quantitative traits being selecting in 75% of the cases (6/8) whereas in binary traits, Relu function was the optimal choice in 63% of the cases (5/8). Moreover, Adam optimizer was the most frequently chosen in binary traits during the hypertuning with 63%. Nevertheless, RMSprop was the optimal option with percentage of 50% in quantitative traits. DL models can capture interactions of large orders because of the presence of hidden layers <u>(Goodfellow et al. 2016</u>, <u>LeCun et al. 2015)</u>. However, RKHS models are also able to capture complex interaction patterns. This ability of both methods can be reflected in our results demonstrating that both can capture complex interactions.

The third aim of this work is to study the impact of various input strategies on the prediction results. It is commonly believed that GP requires a large marker set to be used for an efficient prediction. However, our current results and some of related works <u>(Vourlaki et al. 2022</u>, <u>Bellot et a al. 2018)</u> support that GP models can be effective even with a smaller dataset of markers. However, the optimal marker size can be related to the studied trait <u>(Sandhu et al. 2021)</u>. We also observed that the best input strategy is affected by the chosen phenotypic trait and the training set in some cases (Table 6). Note that MLP models using PCs as input strategy proved beneficial in 66.7% of the cases with MLP as best model. In any case, the different input strategies that we followed indicated that the accommodation of subsets of the markers in GP framework can be equal or even more informative than using the whole marker sets (compared to our results in <u>Vourlaki et al. 2022</u>).

Finally, we would like to mention the challenges and limitations of DL models. Firstly, DL models do not provide clear insights into the genetic architecture of the traits, nor do they give information about the effects of specific markers in the studied traits. Different hyperparameters act on different parts of the data, making it hard to interpret the biological significance and importance of each marker in the model <u>(Bellot et al. 2018</u>, <u>Cuevas et al. 2019)</u>. Also, the high computational cost of training models is a significant drawback, especially when multiple hyperparameters must be optimized for each trait separately <u>(Gulli and Pal et al. 2017)</u>. The outperformance of DL over linear models is not always the case. The prediction ability depends on the studied traits and can be influenced by many factors. There is not a single algorithm that performs better in all species and traits <u>(Perez-Enciso and Zingaretti, 2019)</u> since its performance depends on various factors. Nevertheless, even though the advantage of DL networks against linear methods has not been established yet, their incorporation into plant breeding can be important to improve genetic merit for complex traits.

## Acknowledgements

We acknowledge Josep M. Casacuberta for his helpful advice on the manuscript and Alejandro Navarro Martínez for his contribution to the initial work.

## Funding

Ministerio de Ciencia e Innovación grant PID2019-108829RB-I00 to MPE and RC.

Ministerio de Ciencia e Innovación Postdoctoral Fellowship IJC2020-045949-I to RC.

Programa Severo Ochoa para Centros de Excelencia en I+D" 2020-2023 (CEX2019-000902-S)

## Notes

https://github.com/ivourlaki/Deep-Learning-in-rice-for-prediction

### Competing Interest Statement

The authors have declared no competing interest.

