## Supplementary Tables for "Evaluation of Deep Learning for predicting rice traits using structural and single-nucleotide genomic variants"

Best Hyperparameters for each trait

The most frequently selected hyperparameters over four method categories, MLP, CNN, MLP with PCs and MLP with multiple inputs, for each of the studied traits are summarized in Tables 2-5. As tables depict, the optimal number of layers was 1 and 3 in 37.5% of the studied cases, followed by two layers with percentage of 25%. In the case of the activation function, hyperbolic tangent was the dominant in 50% of the summarized cases (8/16). Linear activation function was the second optimal choice with percentage around 37.5% whereas Rectified Linear Unit (Relu) selected in 12.5% of the cases. Among the available optimizers, Root Mean Square Propagation (RMSprop), Stochastic Gradient Descent (SGD) and Adaptive Moment Estimation (Adam), the most selected were Adam and RMSprop with percentage 43,75% each. In the case of dropout rate, the most frequent value to reduce overfitting in the model was 0.15 with 31.25 % followed by 0.05 with 24%. Concerning the number of filters, the two optimal values were 38 and 64 with equal frequency among the cases (50% each). Finally, we observed that for the number of neurons it was harder to point out one value since various numbers seem to be selected under different conditions. Note that in the case of MLP MULTIPLE, there are six hidden independent layers in which the input marker sets were forwarded. However, these do not count on the “No of hidden layers” (Tables 1-4) since this hyperparameter corresponds to the tuning after concatenating the six layers.

**Table 1:** Optimized hyperparameters for culm diameter.

| Hyperparameter | MLP | CNN | MLP PCs | MLP MULTIPLE |
| --- | --- | --- | --- | --- |
| Activation | Linear | Linear | Tanh | Tanh |
| No of hidden layers | 2 | 2 | 1 | 1 |
| No of neurons | (38,8) | (4,16) | (64) | (38) |
| No of filters | - | 64 | - | - |
| Optimizer | Adam | Adam | Adam | RMSprop |
| Dropout rate | 0.2 | 0.05 | 0.15 | 0.15 |
| Regularization | 0.001 | 0.01 | 0.01 | 0.001 |

**Table 2:** Optimized hyperparameters for leaf senescence.

| Hyperparameter | MLP | CNN | MLP PCs | MLP MULTIPLE |
| --- | --- | --- | --- | --- |
| Activation | Linear | Linear | Linear | Relu |
| No of hidden layers | 1 | 3 | 3 | 1 |
| No of neurons | (16) | (4,8,2) | (128,8,8) | (64) |
| No of filters | - | 38 | - | - |
| Optimizer | Adam | Adam | RMSprop | RMSprop |
| Dropout rate | 0.05 | 0.05 | 0.25 | 0.2 |
| Regularization | 0.001 | 0.001 | 0.01 | 0.001 |

**Table 3:** Optimized hyperparameters for grain weight.

| Hyperparameter | MLP | CNN | MLP PCs | MLP MULTIPLE |
| --- | --- | --- | --- | --- |
| Activation | Linear | Tanh | Tanh | Tanh |
| No of hidden layers | 1 | 3 | 2 | 3 |
| No of neurons | (38) | (16,16,16) | (128,16) | (64,8,2) |
| No of filters | - | 38 | - | - |
| Optimizer | Adam | Adam | RMSprop | RMSprop |
| Dropout rate | 0.1 | 0.05 | 0 | 0.25 |
| Regularization | 0.001 | 0.001 | 0.01 | 0.01 |

**Table 4:** Optimized hyperparameters for time to flowering.

| Hyperparameter | MLP | CNN | MLP PCs | MLP MULTIPLE |
| --- | --- | --- | --- | --- |
| Activation | Tanh | Tanh | Tanh | Relu |
| No of hidden layers | 1 | 3 | 3 | 2 |
| No of neurons | (16) | (2,8,16) | (128,2,8) | (128,8) |
| No of filters | - | 64 | - | - |
| Optimizer | SGD | SGD | RMSprop | RMSprop |
| Dropout rate | 0.1 | 0.05 | 0.1 | 0.25 |
| Regularization | 0.01 | 0.001 | 0.01 | 0.01 |
